# A Fully-Automated Senescence Test (FAST) for the high-throughput quantification of senescence-associated markers

**DOI:** 10.1101/2023.12.22.573123

**Authors:** Francesco Neri, Selma N. Takajjart, Chad A. Lerner, Pierre-Yves Desprez, Birgit Schilling, Judith Campisi, Akos A. Gerencser

## Abstract

Cellular senescence is a major driver of aging and age-related diseases. Quantification of senescent cells remains challenging due to the lack of senescence-specific markers and generalist, unbiased methodology. Here, we describe the Fully-Automated Senescence Test (FAST), an image-based method for the high-throughput, single-cell assessment of senescence in cultured cells. FAST quantifies three of the most widely adopted senescence-associated markers for each cell imaged: senescence-associated β-galactosidase activity (SA-β-Gal) using X-Gal, proliferation arrest via lack of 5-ethynyl-2’-deoxyuridine (EdU) incorporation, and enlarged morphology via increased nuclear area. The presented workflow entails microplate image acquisition, image processing, data analysis, and graphing. Standardization was achieved by i) quantifying colorimetric SA-β-Gal via optical density; ii) implementing staining background controls; iii) automating image acquisition, image processing, and data analysis. In addition to the automated threshold-based scoring, a multivariate machine learning approach is provided. We show that FAST accurately quantifies senescence burden and is agnostic to cell type and microscope setup. Moreover, it effectively mitigates false-positive senescence marker staining, a common issue arising from culturing conditions. Using FAST, we compared X-Gal with fluorescent C_12_FDG live-cell SA-β-Gal staining on the single-cell level. We observed only a modest correlation between the two, indicating that those stains are not trivially interchangeable. Finally, we provide proof of concept that our method is suitable for screening compounds that modify senescence burden. This method will be broadly useful to the aging field by enabling rapid, unbiased, and user-friendly quantification of senescence burden in culture, as well as facilitating large-scale experiments that were previously impractical.

## Introduction

Cellular senescence is a complex stress response typically characterized by essentially irreversible cell cycle arrest, altered morphology, increased lysosomal activity, and profound changes in gene expression, including the acquisition of a senescence-associated secretory phenotype (SASP)^1^. This cellular response can be triggered by many different types of stressors, such as telomere dysfunction^2, 3^, direct DNA damage^4^, oncogenic signaling^5–7^, and mitochondrial dysfunction^8^. Cellular senescence phenotypes are highly heterogeneous and dependent on tissue and cell type^9, 10^. Therefore, detection technologies need to allow for a robust adaptability to different specimens.

Cellular senescence has important physiological roles. For instance, the transient presence of senescent cells is beneficial for cancer prevention^6, 7^, embryonic development^11, 12^, and wound healing^13^. In contrast, senescent cells accumulate in aging tissues^14, 15^, which promotes chronic inflammation and increases the risk of age-related diseases^1^. Preclinical studies have demonstrated that targeting senescent cells can mitigate age-related diseases and increase median lifespan^16^. Hence, drugs that selectively eliminate senescent cells (“senolytics”) or dampen their SASP (“senomorphics”) have the potential to improve the treatment and prevention of age-related diseases^1, 17, 18^. Indeed, several human clinical trials are currently underway^18, 19^, some of which have shown promising outcomes^20^. However, a deeper understanding of this Janus-faced stress response is needed to develop safe and effective senescence-targeting therapies that can combat age-related dysfunction and disease^19^.

One major hurdle in studying cellular senescence is the detection and quantification of senescent cells, primarily because there are no senescence-specific markers^1^. Instead, detection relies on using one or more senescence-associated marker(s) – which are not unique to senescent cells. Some of the most widely adopted senescence-associated markers include senescence-associated β-galactosidase activity (SA-β-Gal) and proliferation arrest measurements, such as lack of EdU incorporation^19^. Because SA-β-Gal is a colorogenic stain, not fluorescent, its quantitative analysis is uncommon. Current methods often rely on manual scoring of microscopy images^21^, or use semi-automated, low throughput image analysis workflows that either do not assess multiple markers^22, 23^ or have limited sample processing capabilities^24^. Thus, quantification of cellular senesce is often subjective and time-consuming, lacking standardization, altogether precluding its use in high-content / high-throughput settings.

To overcome these challenges, we developed the Fully-Automated Senescence Test (FAST). This method produces unbiased assessments of SA-β-Gal and EdU staining by 1) calculation of colorimetric SA-β-Gal optical density (OD), which makes the quantification independent of microscope model and settings; 2) leveraging internal background controls, which allow unbiased staining thresholding and scoring with no assumptions on the senescence phenotype; or alternatively, 3) using machine learning (ML, hence ML-FAST) with biological controls for scoring based on the combination of SA-β-Gal, EdU and nucleus size; 4) automating image acquisition, image processing, and data analysis, which enable high-throughput workflows. We implemented FAST in the commercial image analysis software Image Analyst MKII to provide microplate-based automation and in R to provide custom data analysis and graphing, and here we provide a protocol and all required pipelines and R scripts to implement this assay. FAST is agnostic to the microscopy system used, and we provide examples using a Nikon Eclipse Ti-PFS wide field setup and a Zeiss LSM 980 laser scanning confocal microscope. Moreover, FAST simultaneously allows to evaluate cell counts and morphological alterations – a third senescence hallmark – via nuclear area measurements. Hence, FAST serves as a comprehensive, unbiased tool to rapidly assess senescence burden by measuring three key senescence-associated markers.

## Methods

### Cell culture

Primary human lung microvascular endothelial cells (HMVEC-L) were purchased from Lonza (CC-2527). HMVEC-L were cultured in EGM^TM^-2MV Microvascular Endothelial Cell Growth Medium-2 BulletKit^TM^ (Lonza, CC-3202) at 37°C, 14% O_2_, 5% CO_2_. Human lung fibroblasts IMR-90 were purchased from Coriell Institute (I90). IMR-90 cells were cultured in DMEM (Corning, 01-017-CV) supplemented with 10% FBS (R&D Systems, S11550H), 100 units/mL penicillin, and 100 µg/mL streptomycin (R&D Systems, B21210) at 37°C, 3% O_2_, 10% CO_2_. For all experiments performed, both HMVEC-L and IMR-90 were cultured in 96-well microplates appropriate for microscopy imaging (Corning, 3904), with media changes every 2-3 days.

To achieve serum starvation in HMVEC-L (Fig. 3c), cells were washed twice in DPBS containing Ca^2+^ and Mg^2+^ (Gibco, 14040-117) and then cultured for 72 h in low-serum EGM^TM^-2MV medium (0.5% FBS instead of 5.0% FBS). To achieve high confluency conditions (Fig. 3e), cells were seeded at high density (25,000 cells/cm^2^) and further cultured for 7 days before irradiation; non-senescent control cells were also seeded at the same density and cultured for 7 days before analysis.

### Senescence induction

Senescence was induced as previously described^25^. For ionizing-radiation-induced senescence, the cells were irradiated with 15 Gy, and medium change was performed immediately after treatment. Cells were considered senescent after at least 7 days since irradiation, during which medium was regularly changed (every 2-3 days). For doxorubicin-induced senescence, cells were treated with different dilutions of the drug (Millipore Sigma, D1515-10MG), while non-senescent control cells were treated with vehicle only (DMSO, ThermoFisher Scientific, BP231-100). Cells were cultured in doxorubicin/vehicle-containing medium for 24 h, after which two washes were performed with DPBS containing Ca^2+^ and Mg^2+^ (Gibco, 14040-117) before adding regular medium. Cells were further cultured for 6 days before analysis (i.e. 7 days since the beginning of doxorubicin treatment).

### Senolytic Treatment

On the last day before analysis, cells were treated with the senolytic ABT263/Navitoclax (Selleck Chemicals, S1001) at different concentrations for 24 h, while only vehicle (DMSO) was given as mock treatment.

### SA-β-Gal and EdU Staining

A detailed, step-by-step protocol is provided at protocol.io (dx.doi.org/10.17504/protocols.io.kxygx3ypwg8j/v1). Commercially available kits were used to perform SA-β-Gal (Abcam, ab65351) and EdU staining (ThermoFisher Scientific, C10351). Cells in “Staining wells” were given medium containing 2.5 µM EdU for 24 h before fixation. Cells in “Background wells” were instead given medium containing vehicle (DMSO). After 24h, cells were fixed by adding 8% PFA in PBS pre-warmed to 37°C directly to the medium up to a final concentration of 4% PFA and incubated for 15 min at RT. Subsequently, cells were washed twice with PBS, and SA-β-Gal staining was performed.

To stain for SA-β-Gal, fixed cells were given the staining solution mix as recommended by the manufacturer (Abcam, ab65351). However, the X-Gal powder used was separately purchased (Life Technologies, 15520-018). “Staining wells” were given the complete staining solution mix, whereas “Background” wells were given a solution that did not contain X-Gal, but only vehicle (DMSO). Staining was performed overnight at 37°C in an incubator without CO_2_ control. To prevent nonspecific indole crystal formation, empty spaces in between wells of the microplates were filled with PBS, and parafilm was used to seal the microplates before the overnight incubation.

After the overnight incubation, cells were washed twice with PBS to stop the staining. After SA-β-Gal staining, EdU detection was performed. Briefly, cells were permeabilized at room temperature for 15 min with 0.5% Triton X-100 (Millipore Sigma, T9284-500ML) in PBS. After permeabilization, the Click-iT Reaction Cocktail was added as per the user manual, and cells were incubated for 30 min in the dark. After the incubation, cells were washed once with PBS, counterstained with 0.5 μg/ml DAPI in MilliQ water for 30 min at room temperature in the dark, then washed once with MilliQ water. Finally, cells and cell-free wells were covered with PBS and imaged.

### Image Acquisition

Wide-field microscopy was performed on a Nikon Eclipse Ti-PFS fully motorized microscope controlled by NIS Elements AR 5.21 (Nikon, Melville, NY). The setup comprised of a Lambda 10-3 emission filter wheel, a SmartShutter in the brightfield light path, and a 7-channel Lambda 821 (Sutter Instruments, Novato, CA) LED epifluorescence light source with excitation filters on the LEDs, controlled by a PXI 6723 DAQ (NIDAQ; National Instruments) board. Images were acquired by an Andor iXon Life 888 EMCCD camera (Oxford Instruments) using 10 ms exposure times, with a Nikon S Fluor 10× DIC NA=0.5 lens, using the following filter sets (Semrock; excitation – dichroic mirror – emission given as center/bandwidth in nm): for DAPI: using the 385 nm LED 390/40 – 409 – 460/80; for EdU 480 nm LED with 480/17 – 495 – 542/27. For SA-β-GAL an incandescent Koehler illumination was used and a 692/40 “emission” filter. Using the ND-Acquisition feature of Elements, 3×3 tiled images were recorded without overlap or registration, using the full 1024×1024 resolution of the camera (1.3 µm/pixel), and the above-defined 3 channels. The Kohler condenser was carefully focused for each experiment in the center of a well, with aperture diaphragm semi-open. For autofocusing, the Nikon’s Perfect Focus System was used. For each microplate, two acquisitions were run: one to image the wells containing samples (“Staining” and “Background wells”; typically center 60 wells), and another to image wells that did not contain any cells (“Blank wells”; 36 edge wells). In addition to the above data, the average pixel intensity measured with no illumination (dark current) was determined for precise OD calculation below. Data were saved and analyzed as native *.nd2 files.

### Image Acquisition on an Alternative Setup

Confocal microscopic image acquisition was performed on a Zeiss LSM 980 laser scanning confocal microscope. Standard (Smart Setup) settings were used for DAPI and EdU, and SA-β-GAL was recorded using the transmitted light detector and the 639 nm laser. Koehler illumination and tiling were set up as described above for wide-field microscopy. Here a Plan-Apochromat 10× NA=0.45 lens was used and 3.78 s frame time. Optical zoom of 1.0 resulted in 0.83 µm/pixel resolution in 1024×1024 tile frames. For autofocusing the Zeiss Definite Focus 3 was used. Microplate-based acquisition was set up using the AI sample finder feature of the controller software Zen 2.3, placing one tile region into the center of each well, and data were saved and analyzed as native *.czi files.

### Image Processing

Native format image files were opened in Image Analyst MKII (Image Analyst Software, Novato, CA) as a Multi-Dimensional Open Dialog, representing one microplate (or its blank recordings) at a time. Analysis was performed using modified standard and custom pipelines (https://github.com/gerencserlab/FAST). A detailed, step-by-step protocol is provided at dx.doi.org/10.17504/protocols.io.kxygx3ypwg8j/v1. Briefly, “Blank well” reference images were created by the median projection of replicate wells using the “Create BLANK reference image for multiwell plate using median” pipeline. These reference images in conjunction with pixel intensity related to detector dark current were used then by the main pipeline for SA-β-GAL OD calculation below. Next, the file containing all other assay wells, including “Stained wells” and “Background wells” images was opened, and the pipeline “FAST Analysis Pipeline - Basic” was executed in all wells. The output Excel file containing single-cell measurements for the whole microplate was saved, and further data analysis was triggered by executing the “Run FAST.R Shiny App” pipeline.

### Data Analysis

Single-cell measurements were analyzed using a web browser-based R-application, FAST.R (https://github.com/f-neri/FASTR), which we developed using the R Shiny package^24^ (version 1.7.4, using R 4.3.1) and is designed to run locally. No command line or scripting knowledge is required for its installation or use (for a detailed, step-by-step protocol, see protocol.io: dx.doi.org/10.17504/protocols.io.kxygx3ypwg8j/v1). The inputs of FAST.R are the above Excel output file containing the single-cell measurements, and a plate map in *.csv format. The output consists of i) a single-cell data table, containing single-cell measurements integrated with the user-defined metadata; ii) an analysis report table, which contains staining scoring and nuclear area data for each sample imaged; iii) automatically generated graphs, as shown in the figures below, which help users visualize and understand their data. Briefly, the app associates image analysis results with well condition labels and determines the positive staining thresholds, independently for each condition based on the labels. Then, these thresholds are used for generating positive cell counts and percents for each well and condition. The app also provides additional summary information for each well, such as cell counts, quartile values for each staining, and fold changes. The machine learning based senescence classification was implemented with the R package caret^27^ using random forest classifiers. Briefly, single-cell measurements of all senescence markers (SA-β-Gal, EdU, and nuclear area) were pre-processed by centering and scaling, and then the classifier model was trained using a random forest algorithm and fine-tuned by repeated k-fold cross validation method. The trained classifier was used to calculate the percentage of predicted senescent cells in each well not used for training.

### X-Gal and C_12_FDG co-staining

IMR-90 cells were cultured as described above, except they were seeded on Matrigel-coated glass bottom plates (Greiner SensoPlate #655892). For senescence staining, the culture media was removed from each well containing cells (irradiated and non-irradiated controls) and replaced with custom low sodium bicarbonate, clear imaging media (Image Analyst Software, Novato CA) comprised of DMEM, 1% FBS, L-Gln 4 mM, sodium pyruvate 1 mM, and 25 mM glucose, at 37°C. The media was supplemented with Bafilomycin A (1 µM) and the cultures were incubated for 40 min at 37°C in an air incubator (no CO_2_ control). Cells were finally stained with 30 µM C_12_FDG (ThermoFisher Scientific, D2893) prepared in imaging media (with 1 µM Bafilomycin A and 1 µg/mL Hoechst 33342) and incubated for 1.5 h. Live cell imaging was performed on the above specified Nikon Eclipse Ti-PFS microscope. After imaging C_12_FDG and Hoechst fluorescence, cells were immediately fixed for 10 min in 2% PFA and stained for 20 hours with X-Gal as described above. The microplate was subsequently reimaged for both fluorescence (C_12_FDG and Hoechst) and chromogenic X-Gal staining. Analysis was performed using the “FAST Analysis Pipeline - Basic - Modified for Live plus Fixed Merging” pipeline, where an image registration step matching live- and fixed-cell imaged Hoechst fluorescence was added in front of the basic pipeline and a customized version of the FAST.R Shiny app.

### Statistics

Statistical tests employed are exhaustively described in each figure legend. Such statistical test where either performed using GraphPad Prism or the R package “stats” (v 4.4.0).

## Results

### FAST Workflow

The FAST workflow is comprised of four parts: sample preparation, image acquisition, image analysis, and data analysis (Fig. 1) to assess senescence burden in a fully automated process. To demonstrate these, we used senescence-induced and non-senescent primary human lung endothelial cells (HMVEC-L) and fibroblasts (IMR-90) cultured in optical quality 96-well plates. Regardless of the assayed biology, each microplate included wells containing cells stained with DAPI, EdU, and SA-β-Gal (“Staining wells”), or DAPI only (“Background wells”). Moreover, selected wells contained no cells and no stain but PBS only (“Blank wells”) (Fig. 1a).

**Fig. 1.**
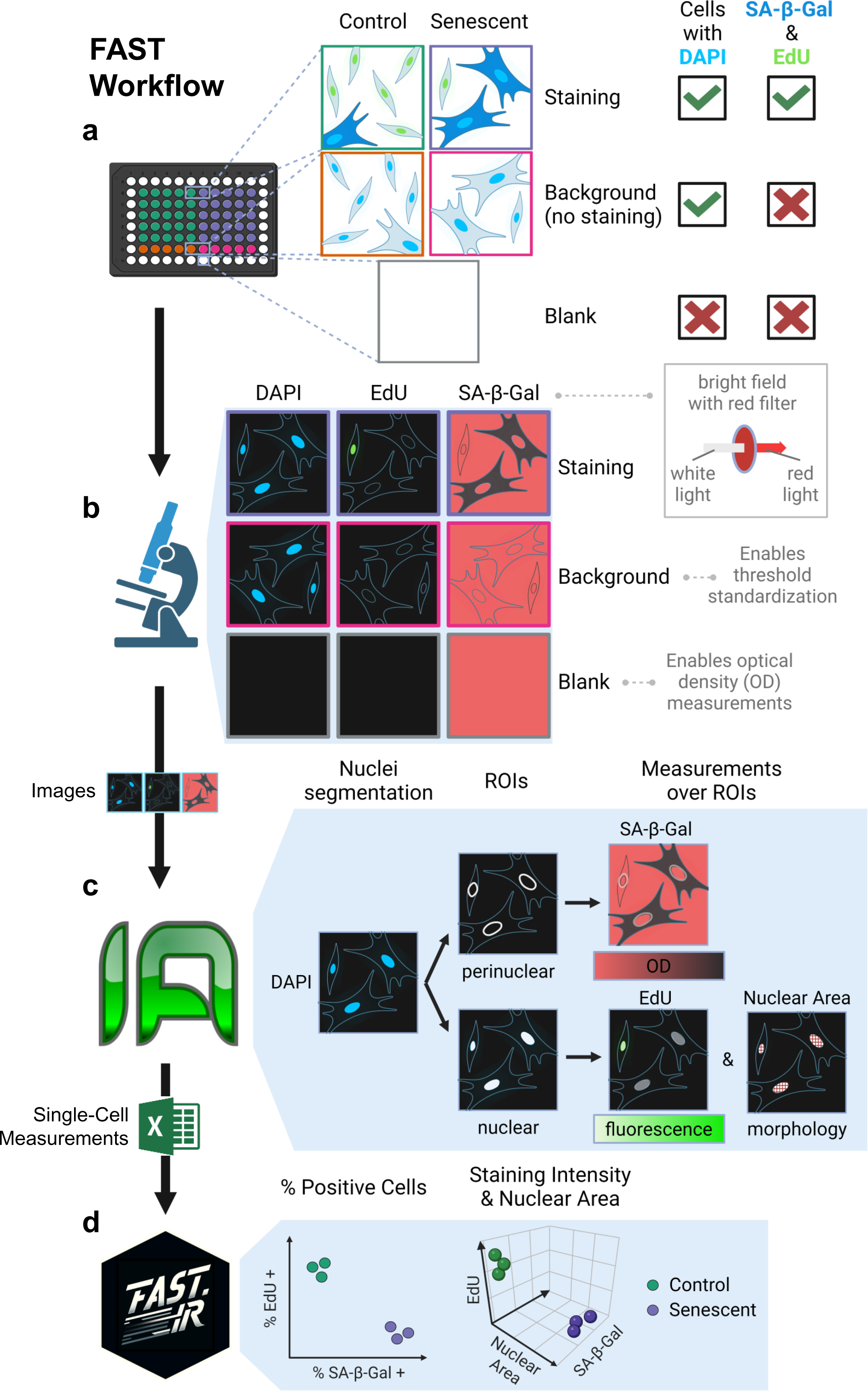
FAST workflow. **a**) For each condition (e.g. Control and Senescent), cells are given substrates for SA-β-Gal and EdU staining (Staining) or vehicle (Background). All cell wells are DAPI-stained. Some wells do not contain any cells (Blank). **b**) Automated image acquisition is performed to capture nuclear counterstain DAPI (blue channel), EdU staining (green channel), and SA-β-Gal (bright field, BF). The use of a red wavelength emission filter (690 nm) for BF imaging results in SA-β-Gal crystals appearing as dark pixels. In combination with the acquisition of Blank images, this modified BF imaging enables optical density (OD) measurements of SA-β-Gal staining. **c**) Image analysis in Image Analyst MKII. DAPI images are segmented, resulting in nuclear and perinuclear labels, and these are used to measure integrated intensities of EdU and integrated ODs of SA-β-Gal, respectively. **d**) Single-cell measurements are analyzed and graphed with FAST.R, a custom R Shiny application.

The entire microplate was imaged, covering most of the bottom of each well using tiling. Fluorescent images were captured for both DAPI (blue channel) and EdU staining (green channel). SA-β-Gal, on the other hand, was imaged as a monochrome bright field channel using a red filter in the light path (or using a red laser to illuminate) matching the peak absorption of the SA-β-Gal staining (Fig. 1b). Image data was saved in a 3-channel *.nd2 or *.czi single file for each microplate, containing a single stitched large view field for each imaged well, and well labels as metadata.

Single-cell EdU intensities, SA-β-Gal optical densities, and nucleus size were determined using automated execution of a single image analysis pipeline for each well in the native microscopy format image data in Image Analyst MKII (Fig. 1c). As a preparation, first, a blank reference image was created from the pixel-wise median of all Blank wells. By calculating the OD using blank images internal to each experiment, optical effects, such as experiment-to-experiment variations in condenser or lamp settings, and vignetting from tiling were canceled. Furthermore, precise OD calculation was obtained by a low-pass spatial filter^25^ to remove non-specific signals in images originating from cellular processes. To generate single-cell data, DAPI images were used to segment nuclei. Here we used a watershed and flood-fill-based morphological segmentation. We chose this method over neural-network-based segmentation, such as Cellpose^26, 27^ (also available in our pipeline repository), because we occasionally observed biases in cell detection due to changing nucleus shape (data not shown). SA- β-Gal absorbance was measured over perinuclear ring-shaped areas and EdU staining intensities over the nuclei. Because in each case the total amount of the marker was relevant to the biology, optical densities or fluorescence intensities were integrated in these areas. The above analysis was performed as a single image analysis pipeline and results were saved as one tabular data (Excel) file per microplate.

For data analysis and visualization, an open-source R-based application, FAST.R, was developed using R Shiny^24^, and was integrated into the above workflow (Fig. 1d). FAST.R allows the association of plate maps with the single-cell data and calculates thresholds to be applied to all “Staining wells” based on the “Background wells”. The application outputs consist of an analysis report, which details staining quartiles, percentages of SA-B-Gal and EdU positive cells, and nuclear area data for each well, as well as graphs presented below. For enhanced senescence detection, senescence scoring can alternatively be performed using a machine learning approach that combines the three staining readouts.

### Automation of the Senescence Marker Scoring

A critical component of automated analysis is the definition of marker positivity using objective criteria. The above workflow provided a standardized input for data analysis by OD calculation for SA-β-Gal and recording of “Background wells” containing unstained cells for both SA-β-Gal and EdU.

We first tested the FAST workflow using primary human lung microvascular endothelial (HMVEC-L) cells induced to senesce through ionizing radiation (IR; Fig 2)^4, 25^. Proliferating cells that were mock irradiated served as the non-senescent control (CTL). The FAST.R app generated detection thresholds for SA-β-Gal and EdU by calculating OD and intensity values at the 95th percentile of cells in a pool of “Background wells” (Fig. 2b.1). Then, these values were used to determine SA-β-Gal and EdU positivity in the “Staining wells” (Fig. 2b.2). Importantly, different conditions, such as IR and CTL, may absorb or scatter light differently, therefore each condition had its own “Background well” control, and FAST.R automatically matched these to the “Staining wells”. Finally, the application computed the percentage of SA-β-Gal- and EdU-positive cells in each well (Fig. 2b.3). As expected, the analysis showed a statistically significant increase in the proportion of SA-β-Gal-positive cells and a concomitant reduction of EdU-positive cells in the IR senescent well compared to non-senescent CTL wells. In addition, FAST.R provided statistics on signal intensities and nuclear area (Fig. 2c-f). Fig. 2 and Supplementary Fig. 1 show graphs automatically generated by FAST.R. We also confirmed that the FAST workflow is compatible with different microscope setups (Supplementary Fig. 2).

**Fig. 2.**
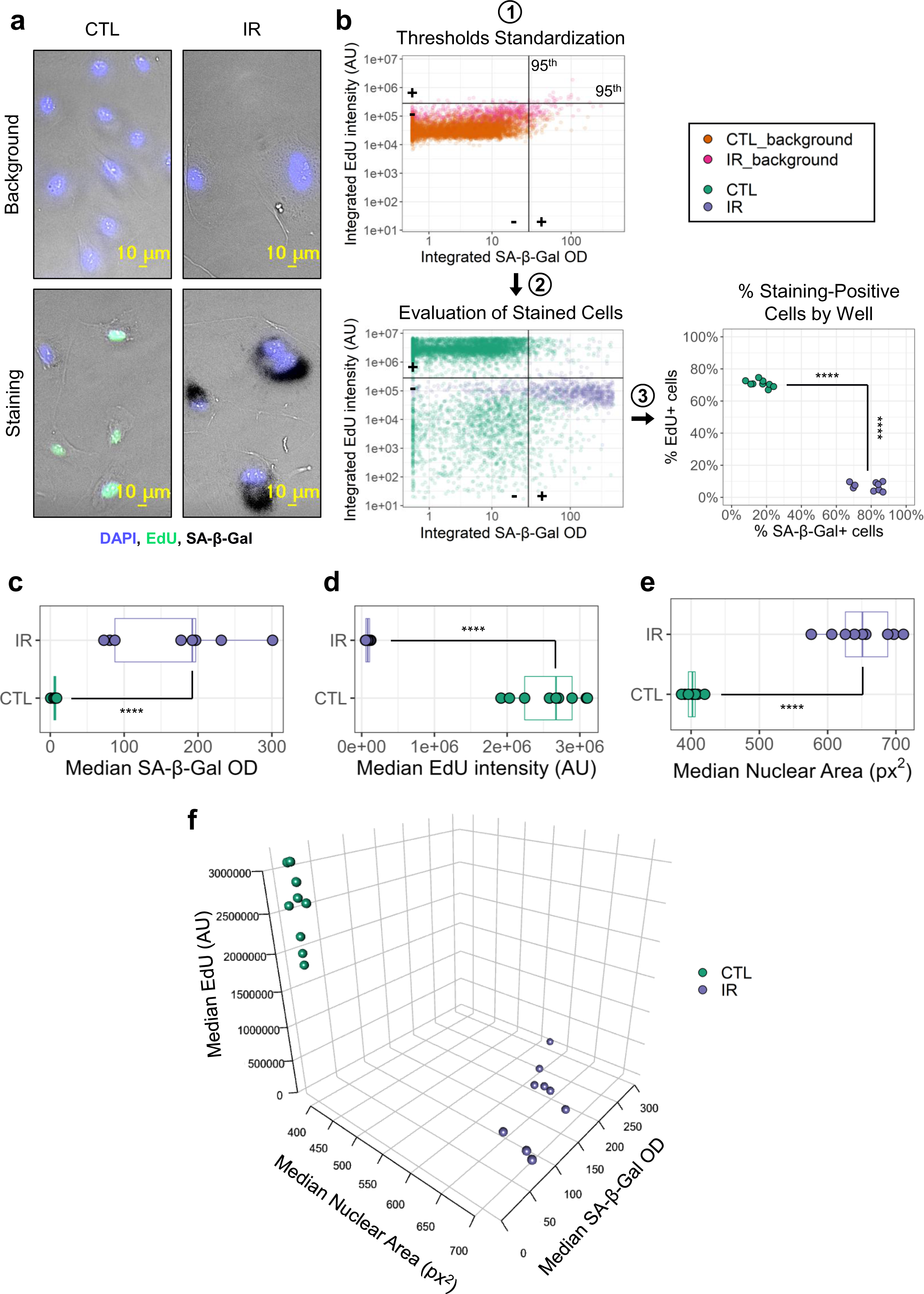
Standardization of detection. **a**) Representative images of primary human microvasculature endothelial cells. Top panels show background cells, while bottom panels show stained cells; left panels show non-senescent cells (CTL), right panels show ionizing radiation-induced senescent cells (IR). **b**) Standardized thresholding for percentage staining calculation. **1**) Signal thresholds are generated based on the 95^th^ percentile in SA-β-Gal and EdU staining measurements of background cells. **2**) Signal thresholds are then used to establish SA-β-Gal and EdU positivity in stained cells. **3**) The percentage of EdU- and SA-β-Gal-positive cells is calculated for each well. **1**-**2**: each dot corresponds to one cell in a representative microplate (n cells: CTL = 6359, CTL background = 5012, IR = 1183, IR background = 884). **3**: each dot corresponds to one well (n = 9) from the same plate. **c-e**) Boxplots with median well values for SA-β-Gal staining (**c**), EdU staining (**d**), and nuclear area (**e**). Each dot corresponds to one well in the same microplate (n = 9). **f**) 3D scatterplot showing all 3 measured variables for each well (n = 9). ****, p<0.0001 by Mann-Whitney test.

### FAST tracks senescent cell populations in experimental settings beset with high rate of false positive staining

The markers SA-β-Gal and lack of EdU incorporation are not specific to cellular senescence. Culturing conditions can result in a significant number of non-senescent cells with false-positive senescence staining, i.e. high SA-β-Gal and low EdU incorporation.

To test the sensitivity of FAST to this intrinsic limitation of senescence-associated markers, we evaluated SA-β-Gal and EdU staining of both senescent and non-senescent cells under conditions known to produce false-positive senescence staining (Fig. 3). Specifically, we analyzed the SA-β-Gal and EdU staining in senescent and non-senescent HMVEC-L cells subjected to serum starvation (Fig. 3a,b), prolonged (48 h) SA-β-Gal staining (Fig. 3c,d), or cultured at a high cell density (Fig. 3e,f). In every experimental condition tested, FAST allowed to detect a statistically significant increase in the proportion of SA-β-Gal positive and EdU-negative cells between the senescent samples and the highly false-positive non-senescent samples (Fig. 3b,d,f).

**Fig. 3.**
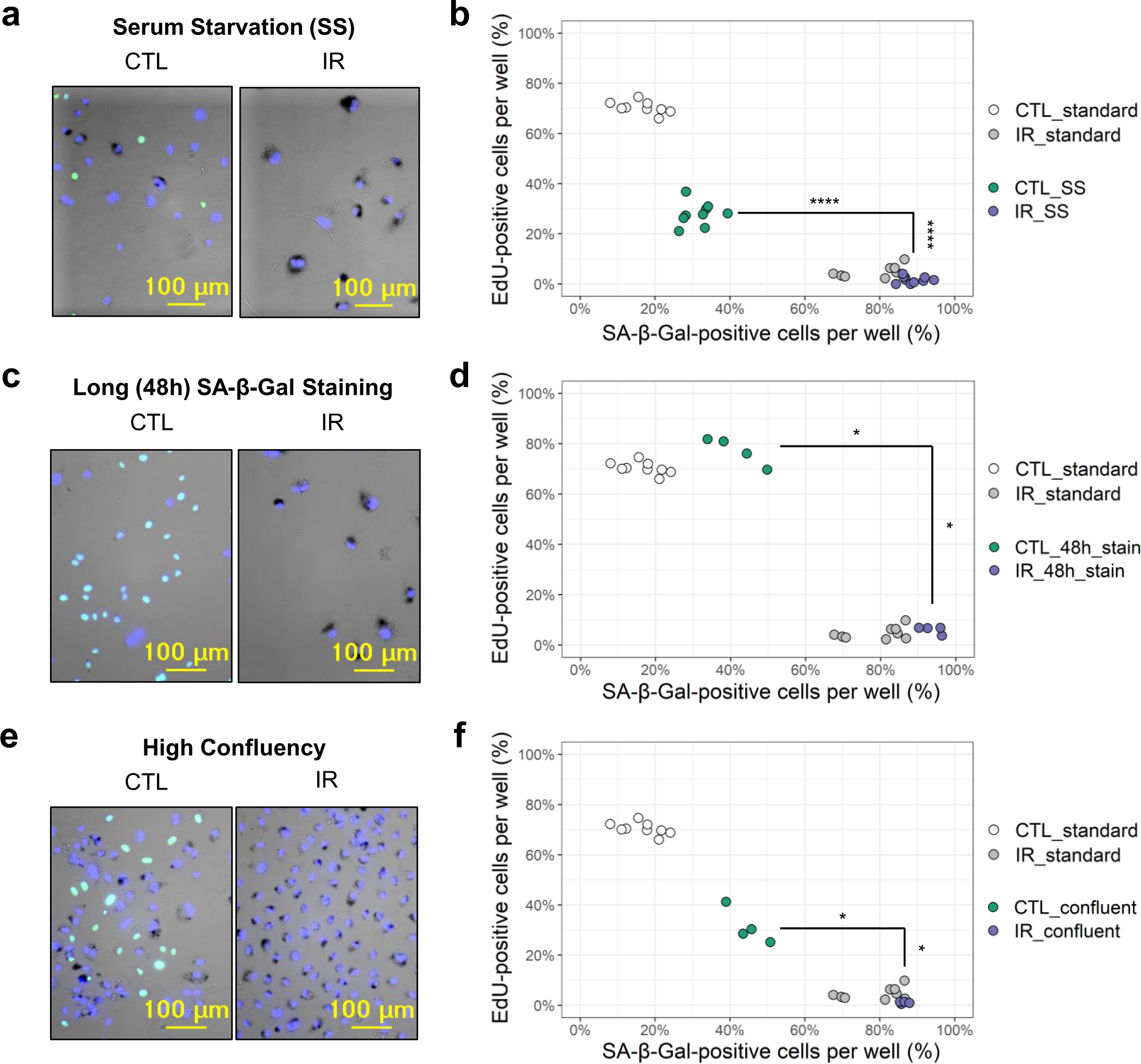
FAST distinguishes senescent cell populations in experimental settings with high rate of false positives. **a**) Representative images of serum-starved (SS) non-senescent (CTL) and ionizing radiation-induced senescent (IR) endothelial cells. **b**) Quantification of (a) with percentages of SA-β-Gal- and EdU-positive cells grouped by well. Both SS and full-serum (standard) samples are shown. Each dot is a well (n = 9) in a representative microplate of 3 independent experiments. **c**) Representative images of CTL and IR endothelial cells with prolonged SA-β-Gal staining (48 h). **d**) Quantification of (c) with percentages of SA-β-Gal- and EdU-positive cells grouped by well. Both samples with prolonged staining (48h stain) and standard overnight staining (standard) are shown. Each dot is a well (n wells: CTL & IR 48h stain = 4, CTL & IR standard = 9). **e**) Representative images of CTL and IR endothelial cells at high confluency. **f**) Quantification of (e) with percentages of SA-β-Gal- and EdU-positive cells grouped by well. Both samples at high confluency (confluent) and low confluency (standard) are shown. Each dot is a well (n wells: CTL & IR confluent = 4, CTL & IR standard = 9). *, p<0.05; ****, p<0.0001 by Mann-Whitney test.

### Comparison of colorimetric X-Gal staining with fluorescent C_12_FDG staining

While FAST relies on the classical SA-β-Gal staining, which uses chromogenic X-Gal as substrate^14^, fluorogenic β-galactosidase substrates have been increasingly employed for cellular senescence detection^31–33^. Despite this, quantitative comparisons of SA-β-Gal staining between X-Gal and fluorogenic dyes are scarce and predominantly restricted to population-level analyses^31^. The principle of the SA-β-Gal staining is detection of an enzyme activity that has been constrained by assay conditions, i.e. suboptimal pH and PFA fixation. Thus, it is unclear how replacing one substrate with another or staining live *vs* fixed cells affects the detection, and therefore whether fluorescence stains are direct substitutes of X-Gal.

Utilizing FAST, here we directly compared the colorimetric and fluorescent SA-β-Gal staining on the single cell level (Fig. 4). To do this, CTL and IR IMR-90 live fibroblasts were incubated with the cell-permeable fluorogenic substrate C_12_FDG (in the presence of 1 µM Bafilomycin A to increase lysosomal pH), followed by live imaging, fixation, staining with X-Gal, and re-imaging the identical view-fields (Fig. 4a). As anticipated, both X-Gal and C_12_FDG staining intensity were elevated in IR-induced senescent cells compared to controls (Fig. 4b). Notably, X-Gal staining exhibited a trending larger relative increase than C_12_FDG (Fig. 4c). Despite both stains being strongly induced by IR, linear regression analysis revealed only a modest correlation at the single-cell level between X-Gal and C_12_FDG staining in IR-induced cultures (R² = 0.142), with no observable correlation in CTL (R² = 0.002) (Fig. 4d). These data question whether the same molecular entity is reported by the two stains. Furthermore, while we successfully combined X-Gal with fluorescence (DAPI, Alexa488), C_12_FDG exhibited redistribution inside and in between cells upon fixation and permeabilization (Supplementary Fig. 3), precluding its use in multiplexed immunocytochemistry paradigms.

**Fig. 4.**
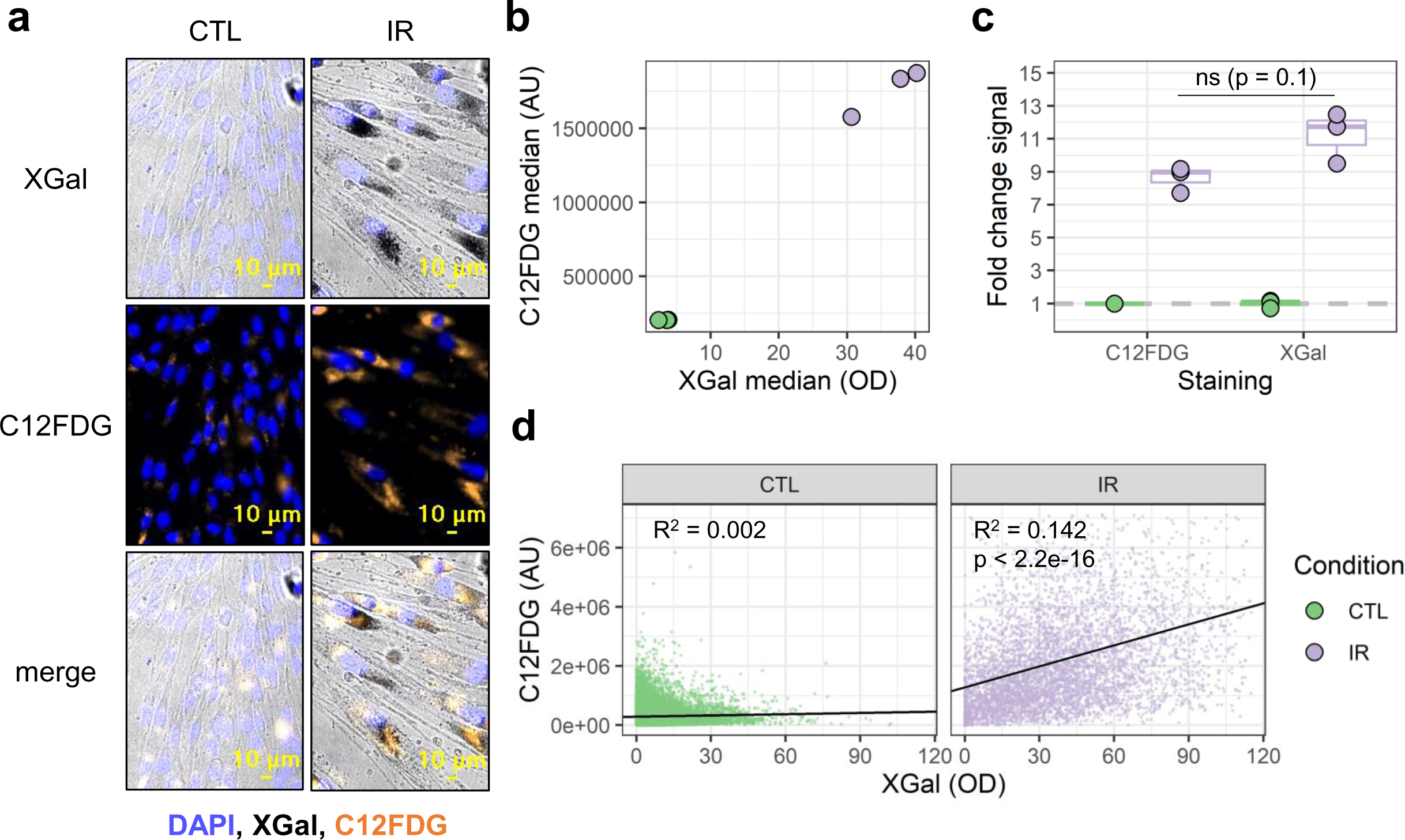
Comparison of colorimetric X-Gal and fluorescent C_12_FDG live cell staining with FAST. **a**) Representative images of non-senescent (CTL) and ionizing radiation-induced senescent (IR) IMR-90 fibroblasts. Live cells were stained with fluorescent C_12_FDG and imaged, followed by fixation, staining with colorimetric X-Gal, and subsequent re-imaging of the same view fields. **b**) Scatterplot with median signal of X-Gal and C_12_FDG for each well (n = 3). **c**) Boxplot showing the fold change in median signal intensity of IR wells relative to CTL (n wells = 3). The p-value was calculated by Mann-Whitney test. **d**) Scatterplots of single-cell staining intensities for X-Gal versus C_12_FDG in CTL (left) and IR (right) conditions. Fitted linear regression models are indicated by solid black lines; n cells: CTL = 22601, IR = 5198. Representative of 3 experiments.

### Benchmarking FAST: Senescence Inducers

The sensitivity of FAST was tested by analyzing dose-responses and calculating Z-factors, in different aspects of cellular senescence. As a proof of concept, first we tested senescence induction in lung fibroblasts IMR-90 after treatment with increasing concentrations of doxorubicin (Doxo), a chemotherapeutic drug known to induce senescence^25, 34, 35^ (Fig. 5a,b). FAST quantifies single cells and it sensitively tracked the expected reduction in cell counts seven days after the commencement of treatment compared to the vehicle-treated (DMSO) cells (Fig. 5c). FAST resolved a concentration-dependent increase in the fraction of SA-β-Gal positive and EdU negative cells (Fig. 5d), consistent with a senescent phenotype, which plateaued at 250 nM and 500 nM for SA-β-Gal and EdU respectively.

**Fig. 5.**
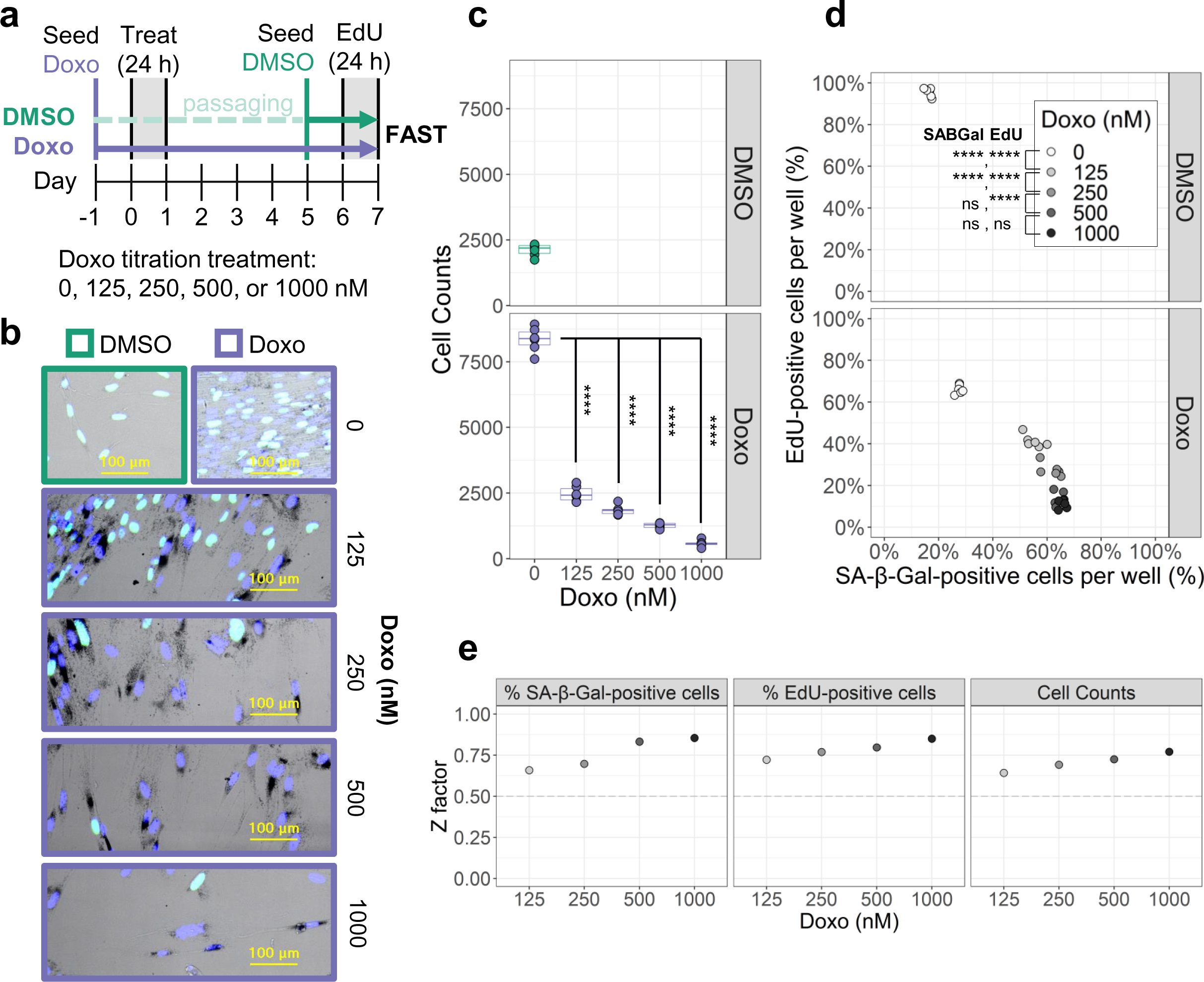
Benchmarking FAST with senescence induction. **a**) Experimental design to test senescence induction with a given compound, in this case, the known senescence inducer doxorubicin (Doxo). **b**) Representative images of primary human IMR-90 fibroblasts treated with different concentrations of Doxo. **c**) Cell counts per well for each Doxo concentration (n wells = 6). **d**) Percentage of SA-β-Gal- and EdU-positive cells per well for each Doxo concentration. Each dot is a well (n = 6). **e**) Z-factor calculations across different metrics: percentage of SA-β-Gal-positive cells, percentage of EdU-positive cells, and cell counts per well. SA-β-Gal and EdU percentages for each Doxo concentration are compared to the DMSO condition. Cell counts for each Doxo concentration are compared to the (8-days cultured) Doxo 0 nM condition. ns, adjusted-p>0.05; ****, adjusted-p<0.0001 by Tukey’s test after significant (p<0.05) one-way ANOVA.

To evaluate the sensitivity of FAST, the Z-factor was calculated (Fig. 5e). The Z-factor is a statistical data quality indicator often used to evaluate the performance and signal robustness of high-throughput screening bioassays ^36^. Assays with Z-factors between 0.5 and 1 are considered to be of good quality, and suitable for high-throughput screenings. Across all the metrics (i.e., SA-β-Gal, EdU, and cell counts), the Z-factor exceeded 0.5 for all concentrations tested. Given that Doxo is a recognized senescence inducer, this data suggests that FAST could serve as a high-quality bioassay for high-throughput screenings of senescence inducers.

### Benchmarking FAST: Senolytics

As a proof of concept, the ability of FAST to measure senolytic activity was tested by treating IR-induced HMVEC-L cells with increasing concentrations of ABT263 (Navitoclax), a chemotherapeutic compound recognized for its senolytic properties^37^ (Fig. 6a). As expected, compared to the non-senescent CTL group (Fig. 6b), the IR-treated HMVEC-L showed an increase in the proportion of SA-β-Gal positive and EdU negative cells. All ABT263 concentrations tested resulted in a significant reduction of viability (based on cell count) in IR cells compared to CTL cells (Fig. 6c). Interestingly, the proportions of SA-β-Gal positive and EdU negative IR cells that survived the senolytic treatment remained the same as that of the vehicle-treated IR cells at all concentrations of ABT263 tested (Fig. 6b and Supplementary Fig. 4).

**Fig. 6.**
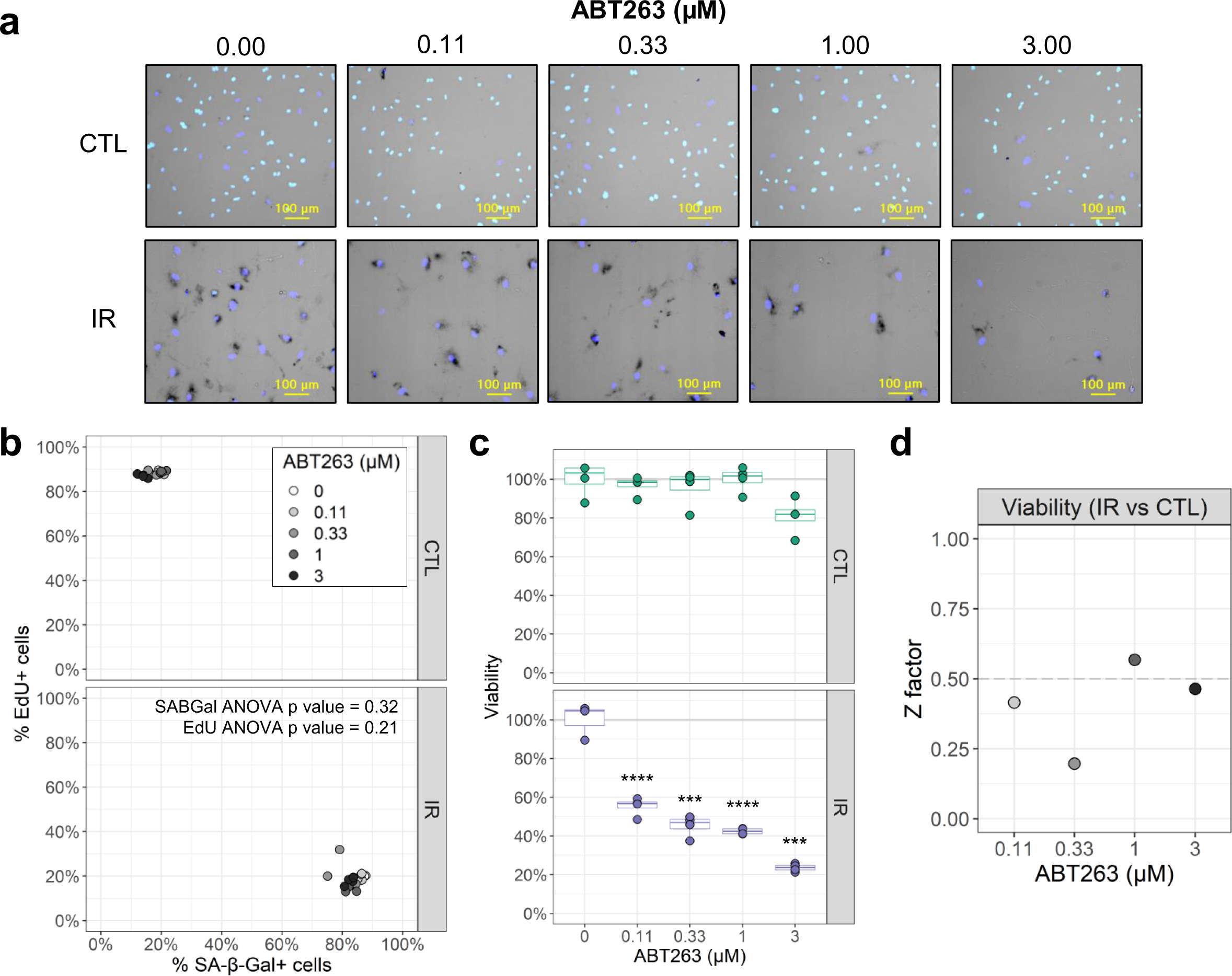
Benchmarking FAST with a senolytic compound. **a**) Representative images of senescent HMVEC-L cells treated with different concentrations of the senolytic compound ABT263 for 24 h. CTL: non-senescent cells, IR: ionizing radiation-induced senescent cells. **b**) Percentage of SA-β-Gal- and EdU-positive cells per well for all ABT263 concentrations (n = 4 wells). **c**) Percentage of viable cells based on cell counts per well normalized to vehicle (0 µM ABT263) condition (n = 4 wells). Comparison of viability between CTL and IR cells for each ABT263 concentration: ***, adjusted-p <0.001; ****, adjusted-p<0.0001 by Bonferroni-Dunn test. **d**) Z-factor calculations for viability measurements at each ABT263 concentration. For each ABT263 concentration, the viability of IR cells was compared to the viability of CTL cells treated with the same senolytic concentration.

To evaluate the sensitivity of FAST at detecting changes in viability upon drug treatment, the Z-factor was calculated (Fig. 6d). One of the ABT263 concentrations tested, 1 µM, had a Z-factor that exceeded 0.5. Considering that ABT263 is a known senolytic, this data suggests that FAST could be used for high-throughput screening of senolytics.

### Analysis of senescence-associated markers using machine learning

The above presented method scores a cell as senescent or non-senescent based on unbiased thresholds for two markers (SA-β-Gal and EdU), determined in unstained samples, thus using no prior knowledge on the senescence phenotype. While nuclear area was informative (Fig. 2e and f), the above threshold generation method of using unstained controls is not applicable for geometric parameters. Therefore, in order to combine all three markers (SA-β-Gal, EdU, and nuclear area), here we describe a trainable, ML approach (ML-FAST) using the unscored, raw data the FAST workflow provides. Classifier models distinguishing senescent from non-senescent cells were based on the random forest algorithm and trained on positive (IR) and negative (mock) sample wells (ignoring impurity) using one, two or all three markers (Fig 7a). The training was performed on single cells (1461-6042 cells) pooled from 6 positive and 6 negative control wells. Because the random forest model requires relatively little training data and time, we performed the training and testing independently for each cell type in each replicate microplate. Fig. 7b and d show model predictions in test wells for HMVEC-L and IMR-90 cells, respectively. The model trained on all three markers was the only one providing consistently greater than 0.5 z-factors in all trials (that includes both cell types, in contrast to Fig. 5), and for this model in all cell conditions, mean z-factors were significantly or trending greater than with all the other models (Fig 7c and e). The combination of nuclear area with SA-β-Gal performed similarly well as combination of EdU with SA-β-Gal, showing small difference to the triple marker model. This latter finding suggest a low-cost variant of FAST that uses only SA-β-Gal and DAPI staining, but no EdU. Altogether, ML-FAST with two or three markers showed high performance across different cell types (HMVEC-L and IMR-90) and culturing conditions (full-serum and serum-starved conditions) as measured by Z-factor (Fig. 7c,e). Thus, the ML-FAST can support sensitive detection of modulators of cellular senescence at a high throughput.

**Fig. 7.**
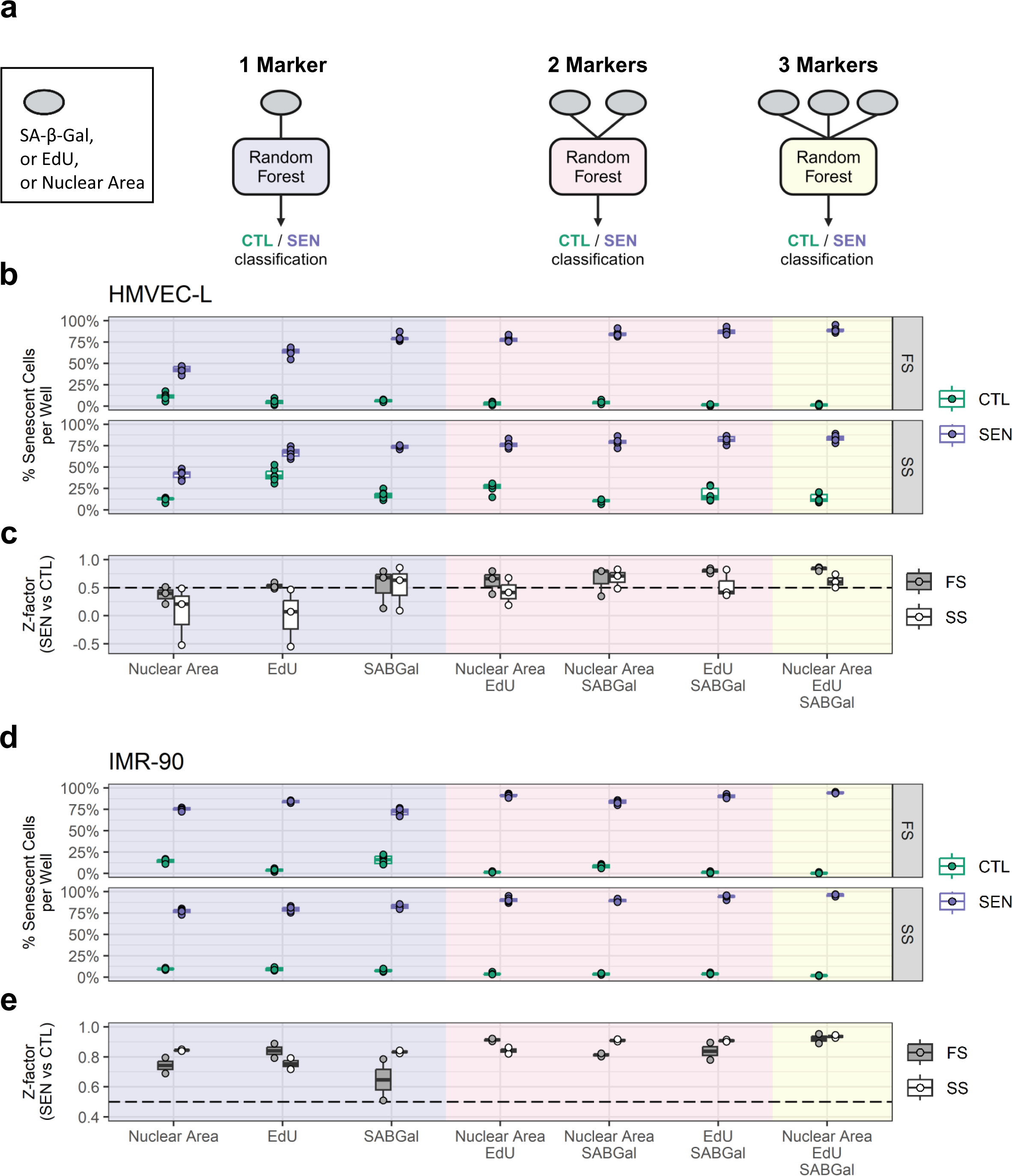
Combination of markers using machine learning improves repeatability of senescence detection. **a**) Conceptual scheme for machine learning classifiers. In each replicate microplate 6 non-senescent (CTL) and 6 IR senescent wells (SEN) including both serum conditions were randomly selected for model training, and the remaining wells were used for the data shown. For each training dataset, random forest classifiers were trained using the indicated combinations of the measured senescence markers (SA-β-Gal, EdU, and nuclear area). **b**) Percentage of cells classified as senescent in CTL (green) and SEN (purple) test wells by each model in HMVEC-L endothelial cells in either full-serum (FS) or serum-starved conditions (SS). Representative of 3 experiments, dots are technical replicate wells (n = 6). **c**) Z-factors comparing SEN and CTL in (b) for both FS (grey) and SS conditions (white). Dots are independent experiments (n=3). **d, e**) Senescence classification in IMR-90 fibroblasts as described (b) and (c), using n = 6 well replicates in d) and n=2 experimental replicates in e). *, p<0.05 using two-way repeated-measures ANOVA with Dunnett’s multiple comparison test comparing to the triple marker classifier. For this all data from c) and e) were pooled and accounted for repeated use of the same data with multiple classifiers. While cell type and serum condition did not have a significant effect, the z-factors significantly varied between the used ML models.

## Discussion

FAST represents a significant advancement over the current manual^19^ and semi-automatic scoring^20–22, 35–37^ methods used to quantify colorimetric SA-β-Gal and EdU staining, which remain low-throughput and not standardized. Key innovations presented here are: 1) calculating SA-β-Gal optical density (OD), which makes the quantification independent of microscope model and settings; 2) implementing internal negative controls (i.e. “Background wells”) to standardize staining thresholding and scoring; 3) automating image acquisition, image processing and data analysis and visualization; 4) combining multiple senescence-associated markers (i.e. SA-β-Gal, proliferation arrest, and enlarged morphology). The pipelines and data analysis app for FAST are readily available, open-source, and support a GUI-based, user-friendly installation and operation for a general biologist audience. A step-by-step protocol is available at protocol.io: dx.doi.org/10.17504/protocols.io.kxygx3ypwg8j/v1).

We implemented image analysis for FAST in Image Analyst MKII because of the microplate-level pipeline-based automation and the fast desktop parallel computing offered by this solution. This software also acts as a GUI to launch all aspects of the analysis. Image Analyst MKII allows manual exploration and well-by-well visualization of cells scored as positive or negative, thus supporting explorative research work. Sequential, unsupervised analysis of multiple microplates is supported by batch-based analysis allowing high throughput applications. Pipelines used here are publicly available and are in human-readable format. Hence, FAST may be also implemented on open-source pipeline-based or scriptable platforms, such as Cell Profiler or FIJI/ImageJ.

Published semi-automated methods^20, 22, 35–37^ fall short of FAST in several of its innovative elements. For example, none of these methods standardized the SA-β-Gal quantification by calculating OD, and, as far as we know, no method employed internal controls to unbiasedly establish staining thresholds, which are instead arbitrarily assigned. Furthermore, the automation component of these workflows was limited. In fact, a few of these relied on manual segmentation^35, 37^, and none of them described and provided protocols for automated sample acquisition, data analysis, or visualization. Finally, often only a single senescence-associated marker (SA-β-Gal) was measured ^20, 35, 37^, and methods measuring more than one marker had limited sample processing capacity^22, 36^.

One limitation of our current work on FAST is that it was only optimized and tested to analyze adherent cell culture samples, unlike other semi-automated, flow cytometry-based workflows that can also analyze ex-vivo samples^22, 36^. However, this limitation could be addressed in future versions of FAST, after adapting it for the analysis of ex-vivo immobilized cell suspensions or tissue cryosections. In fact, the key elements employed by FAST (OD SA-β-Gal measurement, background staining controls, automation, and use of multiple senescence markers combined via ML) remain applicable. In tissue sections, color deconvolution approaches^38^ can be used to convert a color image into OD values specific to stains. A confounder of accurate determination of SA-β-Gal positivity using FAST is that cellular processes and thicker cell bodies can cause a non-specific increase in measured cellular OD not due to light scatter. We mitigated these effects by 1) calculating staining threshold values for SA-β-Gal (and EdU) scoring in a condition-specific manner, thus in control wells with similar cell morphology; 2) applying a low-pass spatial filter during image processing to suppress signal from thin cellular processes. An additional way of suppressing this non-specific signal is increasing the refraction index of the medium during imaging with media such as Optiprep, which has already been used for analyzing formazan OD for succinate dehydrogenase activity cytochemistry^39^.

The ML-FAST approach relies on a classifier trained for the particular experiment using on-plate positive and negative controls, and therefore it is expected to adapt to a broad range of biology. The random forest classifier can be trivially extended using additional markers. The overall cell size is also enlarged in cellular senescence^1^, along with other morphological changes^43–45^, and therefore these can contribute to the identification of senescence phenotypes. However, the visible cell area is also highly dependent on the growth surface available for each cell, therefore on cell density, and this may inadvertently lead to biases. Altogether, ML-FAST can serve as a platform for building more complex cellular senescence assays.

We showed that FAST is capable of distinguishing senescent cells in experimental conditions that cause false-positive senescence staining, due to its graded, quantitative response to staining intensity, thus reducing the intrinsic limitations of the senescence-associated markers used here. We employed FAST across multiple types primary human cells – that is, endothelial cells (Fig. 2,3,6), fibroblasts (Fig 4,5), and astrocytes (data not shown) – and two different senescence-inducing stimuli – specifically, IR (Fig. 1-4,6) and Doxo (Fig. 5). This suggests that FAST can likely be used regardless of the cell type or senescence inducing stimulus used. Moreover, we also show that the FAST workflow is compatible with different automated microscopes (Supplementary Fig. 2).

To the best of our knowledge, we showed for the first time a single-cell-level direct comparison of colorimetric (X-Gal) and fluorescent (C_12_FDG) SA-β-Gal staining. These data confirm the utility of both X-Gal and C_12_FDG in detecting SA-β-Gal activity on the population level. However, they also quantitatively demonstrate that these methods are not interchangeable on the cellular, and perhaps molecular level due to the weak correlation observed. Because C_12_FDG fluorescence was captured before X-Gal staining, this latter could not optically interfere with measuring fluorescence intensity. While X-Gal staining is commonly thought to be incompatible with fluorescence (except for few demonstrated applications^19, 22^) due to its absorption and different microscopy modality, here we showed that X-Gal is better suitable for multiplexed fluorescence assays than C_12_FDG because of the redistribution of the latter during subsequent staining. The X-Gal staining OD values observed at 692nm, near to its absorption peak, were in the range of 0.1-0.2. Below 550nm, its absorption is less than ∼1/5^th^ of the peak^40^ and this equates to absorption of up to 4-8% green fluorescence signal, thus causing little interference. FAST quantifies Alexa488-tagged EdU over the nuclei that typically lack X-Gal staining, further diminishing the possibility of optical interference between probes in this paradigm. For this reason, FAST is especially suitable for combination with other fluorescence stains, especially over the nucleus, such as fluorescence in situ hybridization (FISH) or immunocytochemistry (ICC). The staining protocol for X-Gal is compatible with common FISH and ICC protocols, and analysis of such stains, including spot counting, can be added to the image analysis pipeline and FAST.R application presented here.

Interestingly, we observed that the IR endothelial cells that survived the ABT263 senolytic treatment showed the same fraction of senescence-associated marker positivity as the untreated IR cells (Fig. 6b and Supplementary Fig. 4). We know this is not due to a lack of senolytic activity, as we confirmed that ABT263 preferentially eliminated IR cells compared to control mock-IR cells (Fig. 6c). Thus, ABT263 does not seem to be preferentially targeting IR senescent endothelial cells with canonical senescence staining (SA-β-Gal +, EDU -) compared to senescent cells with non-canonical staining. This could be due to a lack of correlation in IR senescent endothelial cells between senescence-associated staining and expression of ABT263 targets (anti-apoptotic Bcl-2 family proteins). Alternatively, ABT263 might be preferentially targeting the IR senescent cells with canonical senescence staining (SA-β-Gal +, EDU -), but their death might cause a cell non-autonomous cytotoxic effect on the other senescent cells as well. This observation exemplifies how quantitative assaying could propel future studies on mechanisms of cellular senescence and senolysis.

Benchmarks indicate that FAST is suitable for high-throughput screening. Possible applications include identification of environmental pollutants that might exacerbate senescence burden^20, 41^, chemical compound screens for senolytics or compounds that prevent senescence induction, or validation and optimization of senescence induction methods. Because FAST directly measures cell viability while assessing multiple senescence-associated markers, this single assay provides important controls and counter-screen data that can help the identification of compounds with a specific action. In summary, FAST has the potential to substantially advance senescence research by offering a rapid, unbiased, and robust means to assess senescence burden at the single-cell level.

## Acknowledgments

This work was supported by the National Institutes of Health under award numbers U54 AG075932 (Principal Investigators: Campisi and Schilling) and U01 AG060906 (Principal Investigator: Schilling), P01AG017242, P01AG066591, RF1AG068908 (Principal Investigator: Campisi). Figure 1 was generated using Biorender.com.

## Author contributions

F.N. and A.A.G. devised and optimized the method. F.N., S.K.T., C.A.L., and A.A.G. performed the experiments and data analysis. A.A.G. developed Image Analyst MKII and analysis pipelines, and F.N. developed FAST.R. F.N. wrote the initial draft, while A.A.G., P.-Y.D., B.S., and J.C. reviewed and edited the manuscript.

---

**Supplementary Fig. 1.**
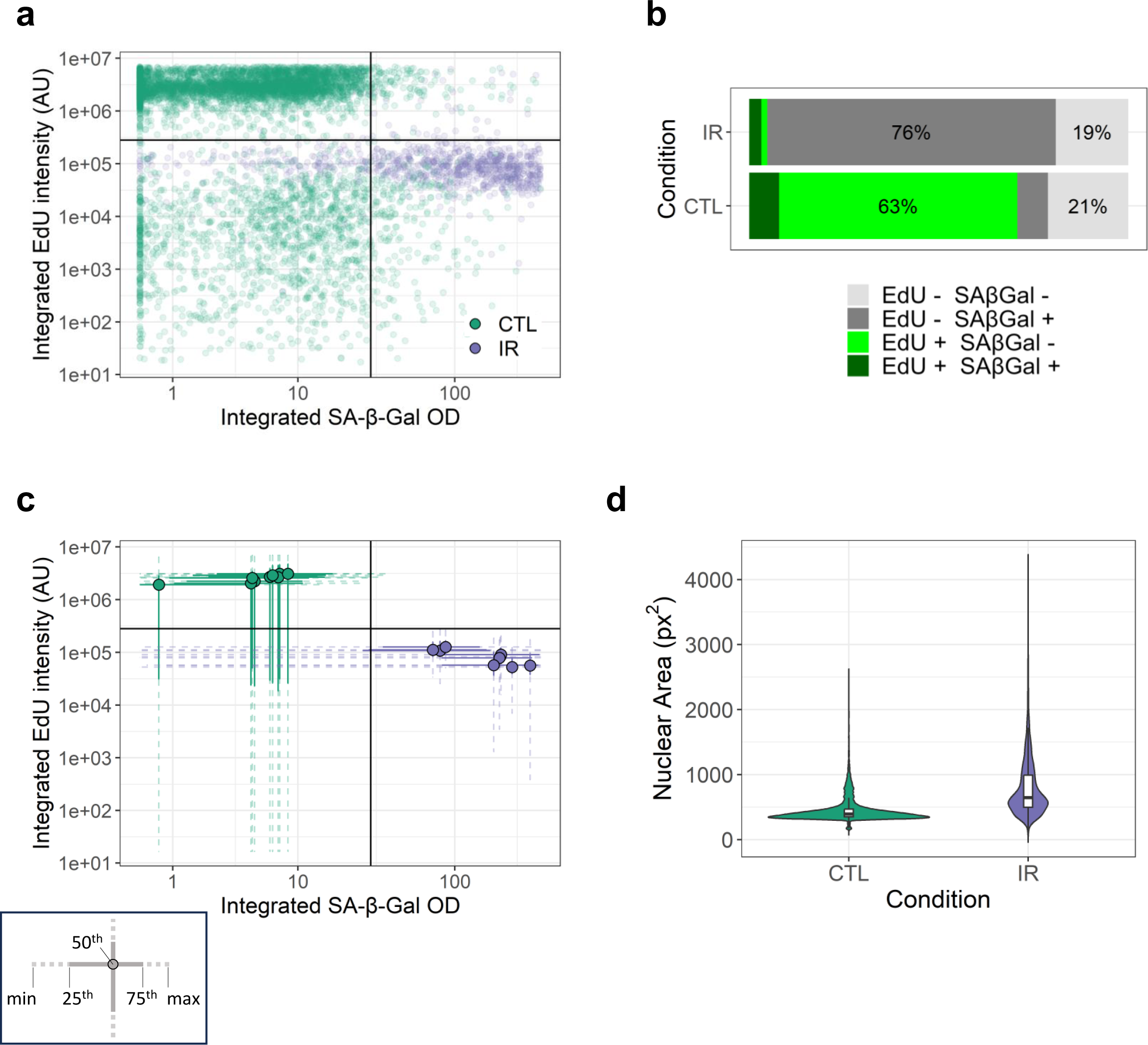
Additional graphs generated with FAST. **a**) Scatterplot showing SA-β-Gal and EdU signal intensity of all cells. Each dot is a cell (n cells: CTL = 6359, IR = 1183). **b**) Bar graph showing the percentage of all cells belonging to one of the four possible staining categories: EdU+/-, SA-β-Gal +/-. **c**) 2D boxplot showing SA- β-Gal and EdU signal intensity of cells grouped by well. Dots indicate median (50^th^ percentile) values, solid lines show interquartile (25^th^ to 75^th^ percentile) range, dashed lines show min to max range. Each data point is a well (n = 9) from the same plate. **d**) Violin plot showing nuclear area distribution.

**Supplementary Fig. 2.**
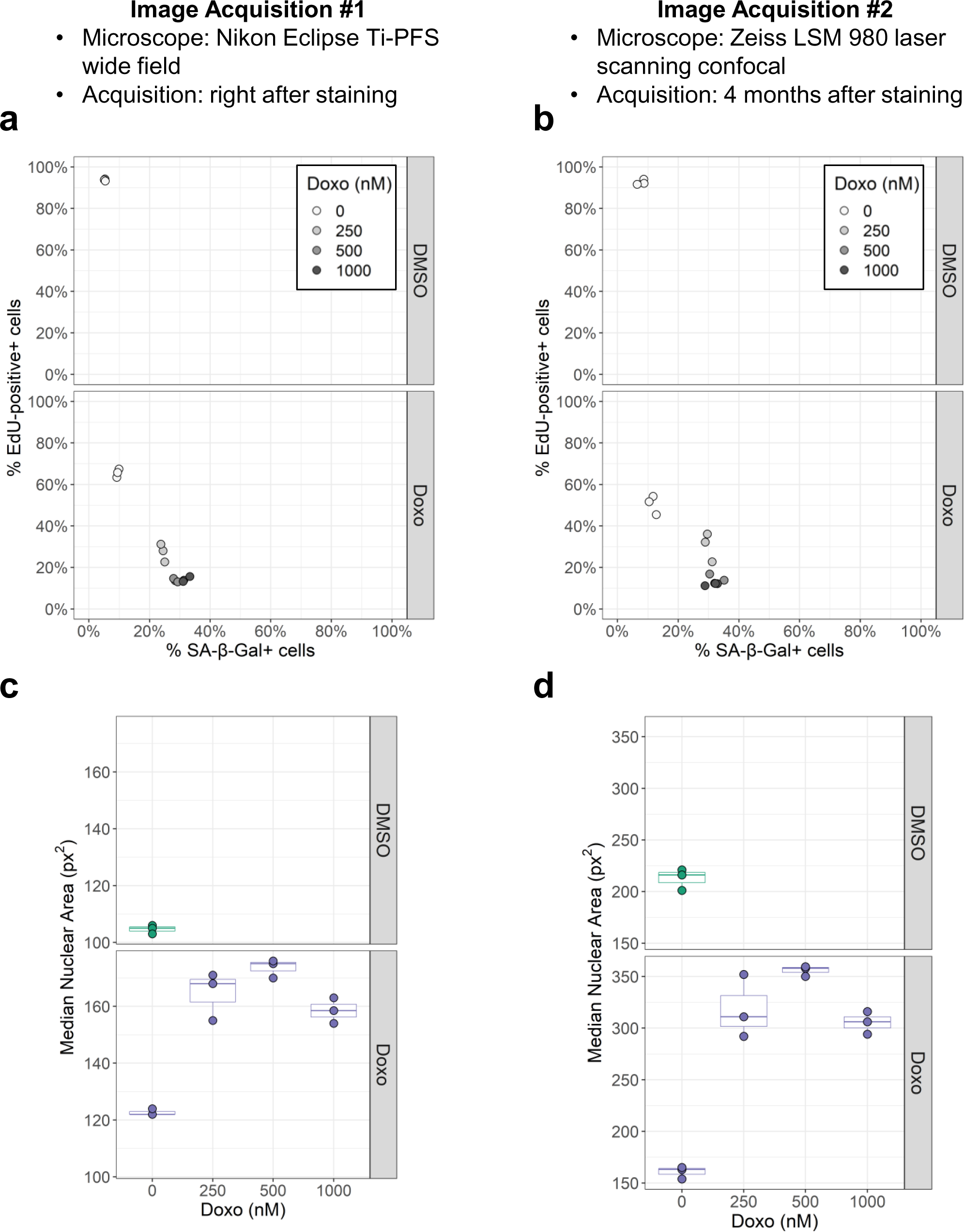
FAST is compatible with different microscope setups. The same microplate containing cells treated with different concentrations of doxorubicin or DMSO vehicle was imaged with a Nikon Eclipse Ti-PFS wide-field microscope (**a**,**c**) and a Zeiss LSM 980 laser scanning confocal microscope (**b**,**d**). **a**,**b**) Percentage of SA-β-Gal- and EdU-positive cells per well for each condition. Each dot is a well (n = 4). **c**,**d**) Boxplot plot showing median nuclear area values for each condition. Each dot is a well (n = 4).

**Supplementary Fig. 3.**
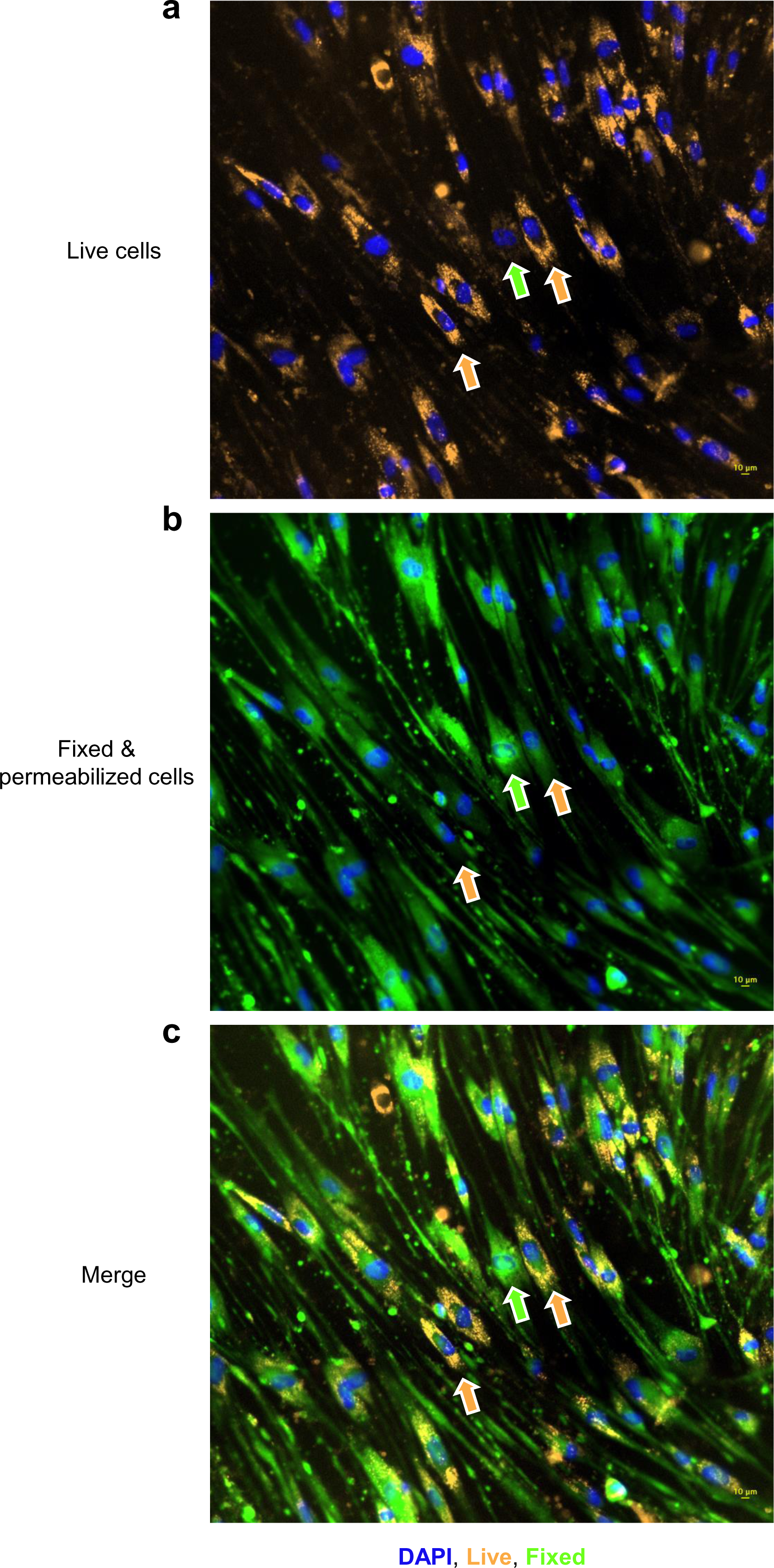
C_12_FDG SA-β-Gal redistributes inside and in between cells upon fixation and permeabilization. **a-c**) Live (a), fixed (b), and merged images (c) of senescent IMR-90 fibroblasts stained with C_12_FDG. For ease of distinction, fluorescence from live images is shown in orange, while fluorescence from fixed cells is shown in green. Orange arrows indicate example cells with bright staining during live imaging subsequently lost after fixation and permeabilization. Green arrow shows an example cell with low staining during live imaging which subsequently becomes highly fluorescent after fixation and permeabilization.

**Supplementary Fig. 4.**
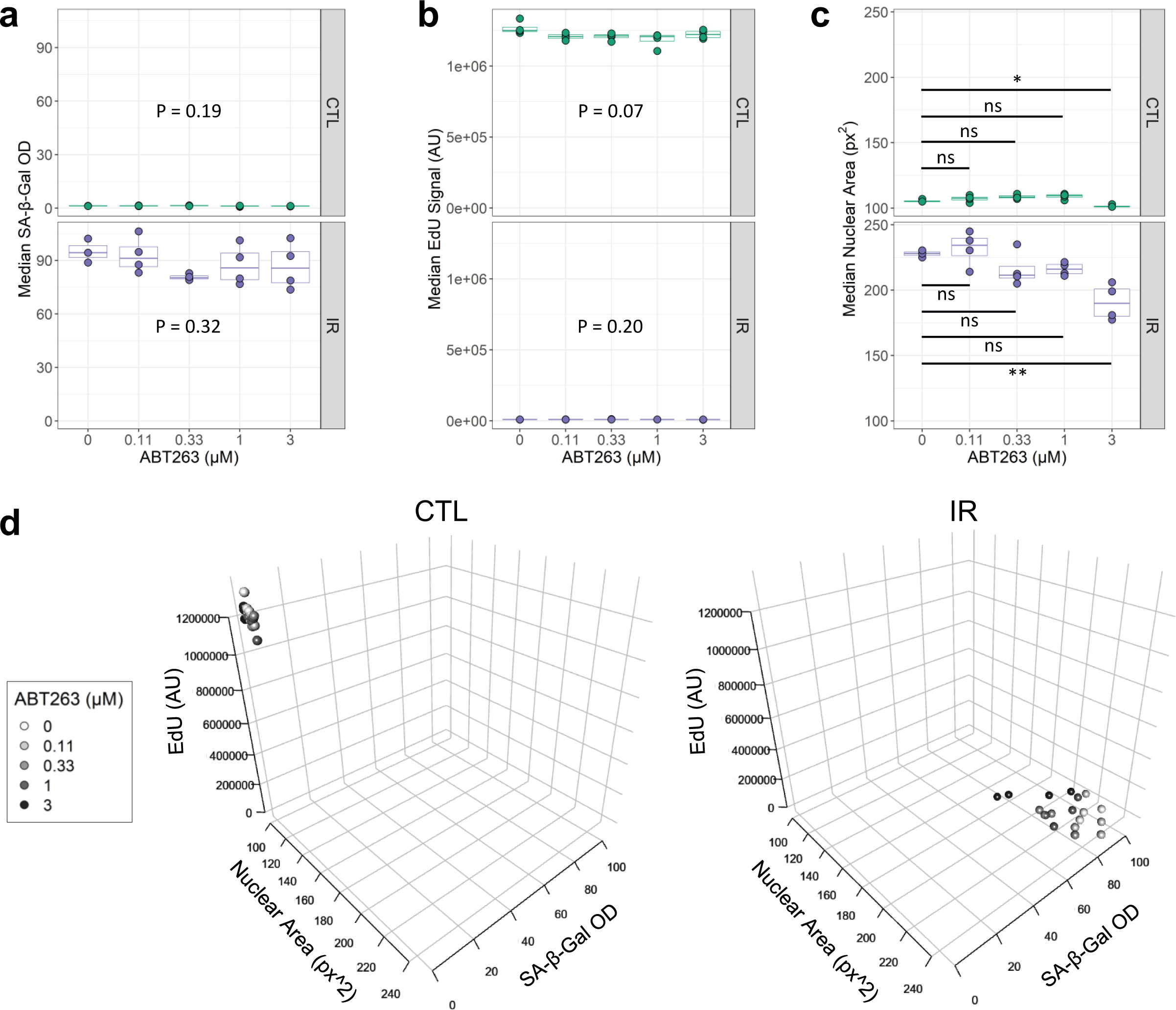
ABT263 treatment does not affect SA-β-Gal and EdU staining in senescent microvascular endothelial cells. **a-c**) Median SA-β-Gal (a), EdU (b), and nuclear area values (c) per well at different ABT263 concentrations in non-senescent control (CTL) and IR-induced senescent (IR) cell populations (n = 4). Non-significant ANOVA p-values (p>0.05) are shown (a,b). ns, adjusted-p>0.05; *, adjusted-p<0.05, **, adjusted-p<0.01 by Tukey’s test after significant (p<0.05) one-way ANOVA. d) 3D scatterplots with all 3 variables for CTL (left) and IR (right) wells at different ABT263 concentrations.

## References

1. Gorgoulis, V. et al. Cellular Senescence: Defining a Path Forward. Cell 179, 813– 827 (2019).

2. Harley, C. B., Futcher, A. B. & Greider, C. W. Telomeres shorten during ageing of human fibroblasts. Nature 345, 458–460 (1990).

3. Bodnar, A. G. et al. Extension of Life-Span by Introduction of Telomerase into Normal Human Cells. Science 279, 349–352 (1998).

4. Leonardo, A. D., Linke, S. P., Clarkin, K. & Wahl, G. M. DNA damage triggers a prolonged p53-dependent G1 arrest and long-term induction of Cip1 in normal human fibroblasts. Genes Dev. 8, 2540–2551 (1994).

5. Serrano, M., Lin, A. W., McCurrach, M. E., Beach, D. & Lowe, S. W. Oncogenic ras Provokes Premature Cell Senescence Associated with Accumulation of p53 and p16INK4a. Cell 88, 593–602 (1997).

6. Di Micco, R. et al. Oncogene-induced senescence is a DNA damage response triggered by DNA hyper-replication. Nature 444, 638–642 (2006).

7. Bartkova, J. et al. Oncogene-induced senescence is part of the tumorigenesis barrier imposed by DNA damage checkpoints. Nature 444, 633–637 (2006).

8. Wiley, C. D. et al. Mitochondrial Dysfunction Induces Senescence with a Distinct Secretory Phenotype. Cell Metab. 23, 303–314 (2016).

9. Cells, S. et al. Unmasking Transcriptional Heterogeneity in Senescent Cells. Curr. Biol. 27, 2652–2660.e4 (2017).

10. Basisty, N. et al. A proteomic atlas of senescence-associated secretomes for aging biomarker development. PLOS Biol. 18, e3000599 (2020).

11. Muñoz-Espín, D. et al. Programmed Cell Senescence during Mammalian Embryonic Development. Cell 155, 1104–1118 (2013).

12. Storer, M. et al. Senescence Is a Developmental Mechanism that Contributes to Embryonic Growth and Patterning. Cell 155, 1119–1130 (2013).

13. Demaria, M. et al. An Essential Role for Senescent Cells in Optimal Wound Healing through Secretion of PDGF-AA. Dev. Cell 31, 722–733 (2014).

14. Dimri, G. P. et al. A biomarker that identifies senescent human cells in culture and in aging skin in vivo. 92, 9363–9367 (1995).

15. Idda, M. L. et al. Survey of senescent cell markers with age in human tissues. Aging 12, 4052–4066 (2020).

16. Xu, M. et al. Senolytics Improve Physical Function and Increase Lifespan in Old Age. Nat. Med. 24, 1246 (2018).

17. Muñoz-Espín, D. & Serrano, M. Cellular senescence: from physiology to pathology. Nat. Rev. Mol. Cell Biol. 15, 482–496 (2014).

18. Di Micco, R., Krizhanovsky, V., Baker, D. & d’Adda di Fagagna, F. Cellular senescence in ageing: from mechanisms to therapeutic opportunities. Nat. Rev. Mol. Cell Biol. 2020 222 22, 75–95 (2020).

19. Chaib, S., Tchkonia, T. & Kirkland, J. L. Cellular senescence and senolytics: the path to the clinic. Nat. Med. 28, 1556–1568 (2022).

20. UNITY Biotechnology Announces Positive 48-Week Results from Phase 2 BEHOLD Study of UBX1325 in Patients with Diabetic Macular Edema | Unity Biotechnology. https://ir.unitybiotechnology.com/news-releases/news-release-details/unity-biotechnology-announces-positive-48-week-results-phase-2/.

21. Kohli, J. et al. Algorithmic assessment of cellular senescence in experimental and clinical specimens. Nat. Protoc. 16, 2471–2498 (2021).

22. Woods, G. & Andersen, J. K. Screening Method for Identifying Toxicants Capable of Inducing Astrocyte Senescence. Toxicol. Sci. 166, 16–24 (2018).

23. Krzystyniak, A., Gluchowska, A., Mosieniak, G. & Sikora, E. Fiji-Based Tool for Rapid and Unbiased Analysis of SA-β-Gal Activity in Cultured Cells. Biomolecules 13, 362 (2023).

24. Biran, A. et al. Quantitative identification of senescent cells in aging and disease. Aging Cell 16, 661–671 (2017).

25. Francesco Neri, Nathan Basisty, Pierre-Yves Desprez, Judith Campisi, and B. S. Quantitative Proteomic Analysis of the Senescence-Associated Secretory Phenotype by Data-Independent Acquisition.

26. Chang, W., et al. shiny: Web Application Framework for R. (2023).

27. Kuhn, M. Building Predictive Models in R Using the caret Package. J. Stat. Softw. 28, 1–26 (2008).

28. Gerencser, A. A., Doczi, J., Töröcsik, B., Bossy-Wetzel, E. & Adam-Vizi, V. Mitochondrial Swelling Measurement In Situ by Optimized Spatial Filtering: Astrocyte-Neuron Differences. Biophys. J. 95, 2583–2598 (2008).

29. Stringer, C., Wang, T., Michaelos, M. & Pachitariu, M. Cellpose: a generalist algorithm for cellular segmentation. Nat. Methods 18, 100–106 (2021).

30. Pachitariu, M. & Stringer, C. Cellpose 2.0: how to train your own model. Nat. Methods 19, 1634–1641 (2022).

31. Kurz, D. J., Decary, S., Hong, Y. & Erusalimsky, J. D. Senescence-associated (beta)-galactosidase reflects an increase in lysosomal mass during replicative ageing of human endothelial cells. J. Cell Sci. 113, 3613–3622 (2000).

32. Martínez-Zamudio, R. I., et al. Senescence-associated β-galactosidase reveals the abundance of senescent CD8+ T cells in aging humans. Aging Cell 20, e13344 (2021).

33. Flor, A. C., Doshi, A. P. & Kron, S. J. Modulation of therapy-induced senescence by reactive lipid aldehydes. Cell Death Discov. 2, 16045 (2016).

34. Maejima, Y., Adachi, S., Ito, H., Hirao, K. & Isobe, M. Induction of premature senescence in cardiomyocytes by doxorubicin as a novel mechanism of myocardial damage. Aging Cell 7, 125–136 (2008).

35. Piegari, E. et al. Doxorubicin induces senescence and impairs function of human cardiac progenitor cells. Basic Res. Cardiol. 108, 334 (2013).

36. Zhang, J.-H. & Oldenburg, K. R. Z-Factor. in Encyclopedia of Cancer (ed. Schwab, M.) 4885–4887 (Springer, Berlin, Heidelberg, 2017). doi:10.1007/978-3-662-46875-3_6298.

37. Chang, J. et al. Clearance of senescent cells by ABT263 rejuvenates aged hematopoietic stem cells in mice. Nat. Med. 22, 78–83 (2016).

38. Shlush, L. I. et al. Quantitative digital in situ senescence-associated β- galactosidase assay. BMC Cell Biol. 12, 16 (2011).

39. Flor, A., Pagacz, J., Thompson, D. & Kron, S. Far-Red Fluorescent Senescence-Associated β-Galactosidase Probe for Identification and Enrichment of Senescent Tumor Cells by Flow Cytometry. JoVE J. Vis. Exp. e64176 (2022) doi:10.3791/64176.

40. Fuhrmann-Stroissnigg, H. et al. SA-β-Galactosidase-Based Screening Assay for the Identification of Senotherapeutic Drugs. JoVE J. Vis. Exp. e58133 (2019) doi:10.3791/58133.

41. Marini, N. et al. Data-driven color augmentation for H&E stained images in computational pathology. J. Pathol. Inform. 14, 100183 (2023).

42. Brand, M. D. et al. Suppressors of Superoxide-H2O2 Production at Site IQ of Mitochondrial Complex I Protect against Stem Cell Hyperplasia and Ischemia-Reperfusion Injury. Cell Metab. 24, 582–592 (2016).

43. Smer-Barreto, V. et al. Discovery of senolytics using machine learning. Nat. Commun. 14, 3445 (2023).

44. Heckenbach, I. et al. Nuclear morphology is a deep learning biomarker of cellular senescence. *Nat*. Aging 1–14 (2022) doi:10.1038/s43587-022-00263-3.

45. Duran, I. et al. Detection of senescence using machine learning algorithms based on nuclear features. Nat. Commun. 15, 1041 (2024).

46. Levitsky, K. L., Toledo-Aral, J. J., López-Barneo, J. & Villadiego, J. Direct confocal acquisition of fluorescence from X-gal staining on thick tissue sections. Sci. Rep. 3, 2937 (2013).

47. Liu, Y. et al. Environmental pollutants exposure: A potential contributor for aging and age-related diseases. Environ. Toxicol. Pharmacol. 83, 103575 (2021).

